# Mechanistic Insights into Dual-Active Liver and Blood-Stage Antiplasmodials

**DOI:** 10.1101/2025.08.06.666330

**Authors:** Mukul Rawat, Nonlawat Boonyalai, Cindy Smidt, Madeline R. Luth, Daisy Chen, Andrew Plater, John Post, De Lin, Joel McMillan, Thomas Eadsforth, Sonia Moliner-Cubel, Oliver Billker, Julian C. Rayner, Francisco-Javier Gamo, Beatriz Baragaña, Elizabeth A. Winzeler, Marcus C. S. Lee

## Abstract

The identification of novel antimalarials with activity against both the liver and blood stages of the parasite lifecycle would have the dual benefit of prophylactic and curative potential. However, one challenge of leveraging chemical hits from phenotypic screens is subsequent target identification. Here, we use *in vitro* evolution of resistance to investigate nine compounds from the Tres Cantos Antimalarial Set (TCAMS) with dual liver and asexual blood stage activity. We succeeded in eliciting resistance to four compounds, yielding mutations in acetyl CoA synthetase (AcAS), cytoplasmic isoleucine tRNA synthetase (cIRS), and protein kinase G (PKG) respectively. Using a combination of CRISPR editing and in vitro activity assays with recombinant proteins, we validate these as targets for TCMDC-125075 (AcAS), TCMDC-124602 (cIRS), and TCMDC-141334 and TCDMC-140674 (PKG). Notably, for the latter two compounds, we obtained a T618I mutation in the gatekeeper residue of PKG, consistent with direct interaction with the active site, which we modelled with molecular docking. Finally, we performed cross-resistance evaluation of the remaining five resistance-refractory compounds using the Antimalarial Resistome Barcode sequencing assay (AReBar), which examined a pool of 52 barcoded lines with mutations covering >30 common modes of action. None of the five compounds where *in vitro* evolution of resistance was not successful yielded validated hits using AReBar, indicating they likely act via novel mechanisms and may be candidates for further exploration.

## Introduction

Malaria continues to pose a major global health challenge, responsible for millions of infections and thousands of deaths annually [1]. Although several antimalarial drugs are currently in use, the growing prevalence of drug resistance underscores the urgent need for new therapeutic strategies. Since 2001, artemisinin-based combination therapies (ACTs) have served as the frontline treatment for malaria [2]. These therapies pair artemisinin, which rapidly reduces parasite levels, with a longer-acting partner drug to ensure full clearance of the infection. However, resistance to artemisinin has been well documented in Southeast Asia and early indications of reduced efficacy have also been reported in parts of Africa [3–6]. In response, a third drug has been introduced in some regions; such triple artemisinin-based combination therapies (TACTs) have shown potential in overcoming declining ACT effectiveness [7, 8]. Despite these efforts, the lengthy process of developing new antimalarial agents highlights the necessity of continued research into novel drug targets and combinations to maintain treatment efficacy and protect vulnerable populations from malaria.

Although many antimalarial compounds have been identified through phenotypic screening, most are effective only against the blood stage of the parasite. However, the identification of targets that also confer vulnerability at other lifecycle stages would have additional advantages in providing prophylactic activity in the case of liver-stage actives, or transmission-blocking activity in the case of gametocyte-stage actives [9–12]. Compounds used for prophylaxis would also benefit from the considerably lower parasite numbers during the liver stage, potentially ameliorating the risk of resistance development.

The Research Centre for Diseases of the Developing World (DDW) at Tres Cantos, GSK, identified 13,533 molecules, known as the Tres Cantos Antimalarial Compound Set (TCAMS), with activity against the blood stages of *Plasmodium* parasites [13]. This compound set was subsequently assessed for activity against the liver-stage using *P. berghei* infected HepG2 cells. This screen identified 103 compounds with dual-stage activity, showing effectiveness against both blood- and liver-stage parasites [14]. However, the mode of action for many compounds remains unclear.

Here we evaluated nine dual-active inhibitors identified in this screen (**Figure 1A**) [14]. We conducted mode of action studies through *in vitro* evolution of drug resistance and whole genome sequencing followed by target validation using *in vitro* assays with genetically modified parasites and biochemical assays with recombinant proteins, identifying the targets for four compounds.

**Figure 1.**
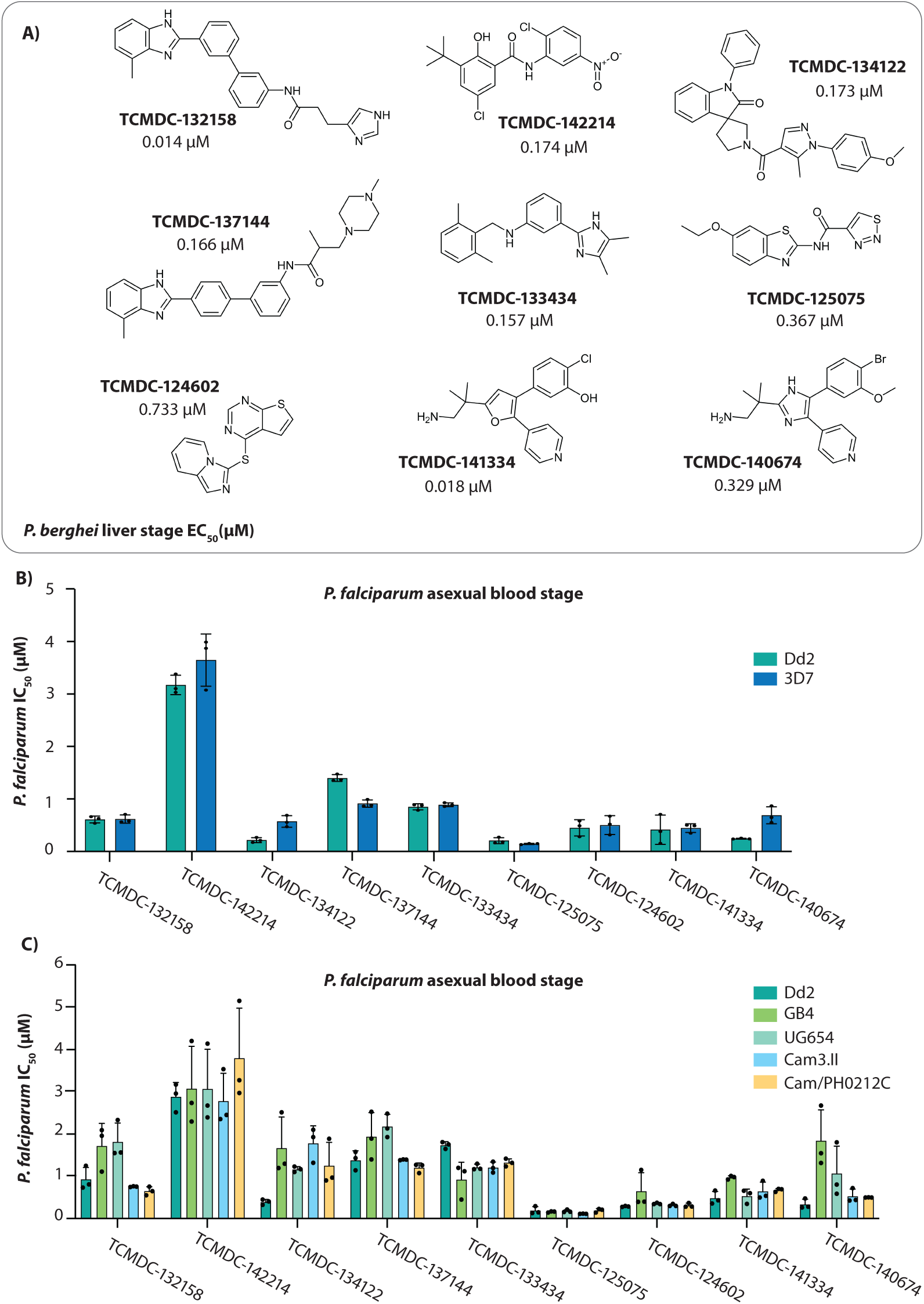
Antiplasmodial activity of selected compounds. **(A)** Chemical structures of nine compounds from the Tres Cantos Antimalarial Compound Set (TCAMS) with dual-stage activity against both blood and liver stages of *Plasmodium*. Liver-stage activity was obtained from **(9)**. **(B)** *In vitro* efficacy of the nine compounds against *P. falciparum* laboratory strains 3D7 and Dd2. **(C)** Antiplasmodial activity of the compounds across a panel of geographically diverse *P. falciparum* strains including Dd2 (Laos), GB4 (Ghana), UG654 (Uganda), Cam3.II and Cam/PH0212C (Cambodia). Activity data for all compounds are detailed in **Supplementary Tables 1 and 2**. In (B) and (C), each dot represents a biological replicate (n = 3); error bars indicate mean ± SD.

## Results

### Activity of liver-stage active compounds against diverse *P. falciparum* strains

Half-maximal growth inhibition concentrations were determined for the nine query compounds against chloroquine-sensitive (3D7) and multidrug-resistant (Dd2) parasites (**Figure 1B, Supplementary Table 1**). Compounds were further assessed against a panel of geographically diverse parasite strains, including two multidrug-resistant Cambodian isolates (resistant to artemisinin, chloroquine, and pyrimethamine) and strains from Uganda (UG659) and Ghana (GB4). These parasite strains displayed similar sensitivity to the tested compounds, with potencies largely comparable to those observed in the laboratory strain Dd2 (**Figure 1C, Supplementary Table 2**). However, a reduction in sensitivity to TCMDC-134122 was noted across all strains when compared to Dd2. Similarly, decreased sensitivity to TCMDC-140674 was observed in strains UG659 and GB4 (**Figure 1C, Supplementary Table 2**).

*In vitro* evolution and whole genome analysis (IVEWGA) is a powerful reverse genetics tool used to identify drug targets [15]. In this method, parasites are exposed to selective concentrations of compound (3×IC_50_), resulting in the enrichment of the parasites carrying mutations in the drug target or a resistance determinant. This is followed by whole genome sequencing to identify candidate mutations responsible for conferring resistance. To aid the process of resistance generation, our lab has previously developed a mutator parasite line (Dd2-Polδ) with a mutation in DNA polymerase δ [16]. The mutations in DNA polymerase δ result in defective proofreading activity of the enzyme, leading to higher mutation rates and, consequently, a higher propensity to generate resistance. Dd2-Polδ parasites were found to generate resistance with a lower minimum inoculum for resistance (MIR) [16].

Resistance selection experiments were performed using a single-step selection protocol with 5×10⁷ Dd2-Polδ parasites performed in triplicate. Recrudescence was observed within 16-25 days for four compounds: TCMDC-141334, TCMDC-140674, TCMDC-125075, and TCMDC-124602 (see **Figures 2A, 3A, 4A and 4C**). To further characterise the recrudescent lines, bulk cultures were first subjected to limiting dilution cloning, and clonal populations were isolated for whole-genome sequencing. Two clones from each selection flask were sent for whole genome sequencing analysis.

**Figure 2.**
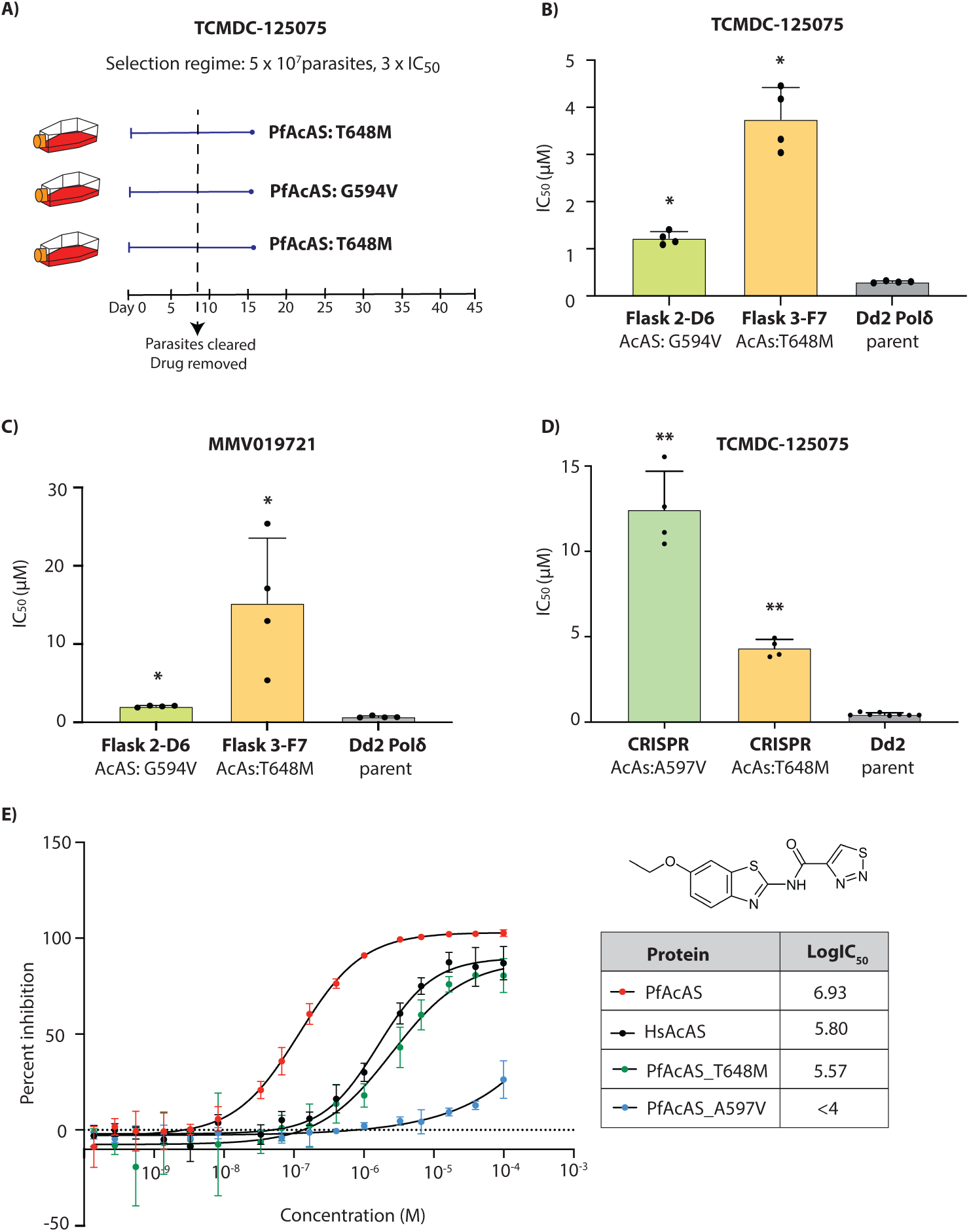
TCMDC-125075 targets *P. falciparum* acetyl-CoA synthetase (*Pf*AcAS). **(A)** Single-step resistance selection against TCMDC-125075 was performed in triplicate using 5 × 10⁷ DNA polymerase-mutator parasites (Dd2-Polδ). All three selections yielded mutations in PfAcAS. **(B)** Clonal lines derived from two independent flasks (Flask 2 and Flask 3) were tested for resistance to TCMDC-125075. The parental Dd2-Polδ line served as a control. **(C)** The same clones (Flask2-D6 and Flask3-F7) were also tested against a structurally related *Pf*AcAS inhibitor, MMV019721. **(D)** CRISPR-edited lines carrying PfAcAS mutations T648M and A597V displayed reduced sensitivity to TCMDC-125075. Dd2 parental line was used as a control. **(E)** *In vitro* enzymatic inhibition assays with recombinant *Pf*AcAS, human AcAS (*Hs*AcAS), and mutant *Pf*AcAS proteins (T648M and A597V) demonstrated selective activity of TCMDC-125075 against PfAcAS. **(B-E)**, each dot represents a biological replicate (n = 4); bars indicate mean ± SD and statistical significance determined by Mann-Whitney *U* tests (*p<0.05, **p<0.01).

For the remaining five compounds, which initially did not yield resistant parasites, selection experiments were repeated using a higher inoculum of 1×10⁸ Dd2-Polδ parasites. Parasite regrowth was observed after 25–30 days for four of these compounds—TCMDC-132158, TCMDC-134122, TCMDC-137144, and TCMDC-133434, while no regrowth was detected for TCMDC-142214. However, dose response assays for the four compounds showing recrudescence did not demonstrate any shift in IC₅₀ values, suggesting a lower potential for resistance development for these compounds.

### Whole genome sequencing of TCMDC-125075 identified a mutation in *P. falciparum* acetyl CoA synthetase (PfAcAS)

Drug sensitivity assays on TCMDC-125075-selected clones revealed a significant increase in IC_50_ (**Figures 2B).** Whole-genome sequencing of 2-3 clones from each of the three independent selection flasks revealed mutations in the gene encoding acetyl CoA synthetase (*Pf*AcAS, PF3D7_0627800; **Supplementary Table 3**). Two distinct mutations in *Pf*AcAS were identified: T648M, present in all clones from Flasks 1 and 3, and G594V, found in both clones from Flask 2 (**Figure 2A**). The G594V mutation was associated with a lower level of resistance (**Figure 2B**). No significant copy number variants (CNVs) were detected in the whole-genome sequencing analysis (**Supplementary Table 4**). Interestingly, the T648M mutation in *Pf*AcAS had been previously observed in resistance studies involving MMV019721 [17]. Since MMV019721 and TCMDC-125075 are structurally related (heterocyclic core linked to aryl or heteroaryl rings), we evaluated TCMDC-125075-resistant parasites against MMV019721, demonstrating cross-resistance (**Figure 2C**). To validate the importance of *Pf*AcAS for resistance to TCMDC-125075, we used previously generated CRISPR-edited mutant lines bearing either a T648M or A597V mutation in *Pf*AcAS [17]. Both CRISPR-edited mutant lines showed significantly reduced susceptibility to TCMDC-125075 with 5-fold and 12.5-fold higher IC_50_ for the *Pf*AcAS-T648M and *Pf*AcAS-A597V lines, respectively (**Figure 2D**). Overall, these assays indicate that mutations in *Pf*AcAS drive resistance to TCMDC-125075.

Next, we demonstrated the ability of TCMDC-125075 to inhibit recombinant *Pf*AcAS using an *in vitro* assay that quantifies the enzymatic product, acetyl-CoA (**Figure 2E**). We also evaluated the compound against human AcAS (*Hs*AcAS), and two *Pf*AcAS mutants with the same polymorphisms as the CRISPR lines tested above: T648M and A597V. The human AcAS enzyme exhibited ∼10-fold reduced sensitivity to TCMDC-125075. Similarly, both mutant *Pf*AcAS enzymes also exhibited higher IC_50_ values compared to the wild-type *Pf*AcAS, indicating reduced sensitivity to the inhibitor (**Figure 2E, Supplementary Table 5**). Consistent with dose–response assays in parasites, the PfAcAS-A597V mutant protein demonstrated the weakest potency to TCMDC-125075 *in vitro* (**Figure 2E, Supplementary Table 5**).

### TCMDC-124602 targets *P. falciparum* cytoplasmic isoleucyl-tRNA synthetase (*Pf*cIRS)

Resistance selections with TCMDC-124602 yielded recrudescent parasites in 3 out of 3 cultures starting with 5×10^7^ parasites of the Dd2-Polδ line **(Figure 3A)**. Previous studies had identified multistage-active thienopyrimidine compounds (MMV1091186, MMV1081413, MMV019869 (TCMDC-124553), and MMV019266 (TCMDC-123835)) as targeting PfcIRS in *Plasmodium* [18]. Our selected compound, TCMDC-124602 (MMV019904), was found to share close structural similarities with these molecules, including common core scaffolds such as a thiazole ring, pyrimidine ring, and a sulphur linker. As expected, whole genome sequencing of TCMDC-124602-resistant clones identified two mutations in *Pf*cIRS: F578C and N268I (**Figure 3A, Supplementary Table 6**). No detectable CNVs were observed in the whole-genome sequencing data (**Supplementary Table 4**). The clone with the F578C mutation (Flask 2) exhibited a relatively small shift in IC_50_ (1.7×IC_50_) compared to the clone in Flask 3 that harboured the N269I mutation, which showed a 9.3×IC_50_ shift (**Figure 3B**). Notably, a different mutation at this amino acid residue 269 (N269K) was previously identified in selections with MMV1091186 [18], indicative of its importance in resistance to this class of compounds.

**Figure 3.**
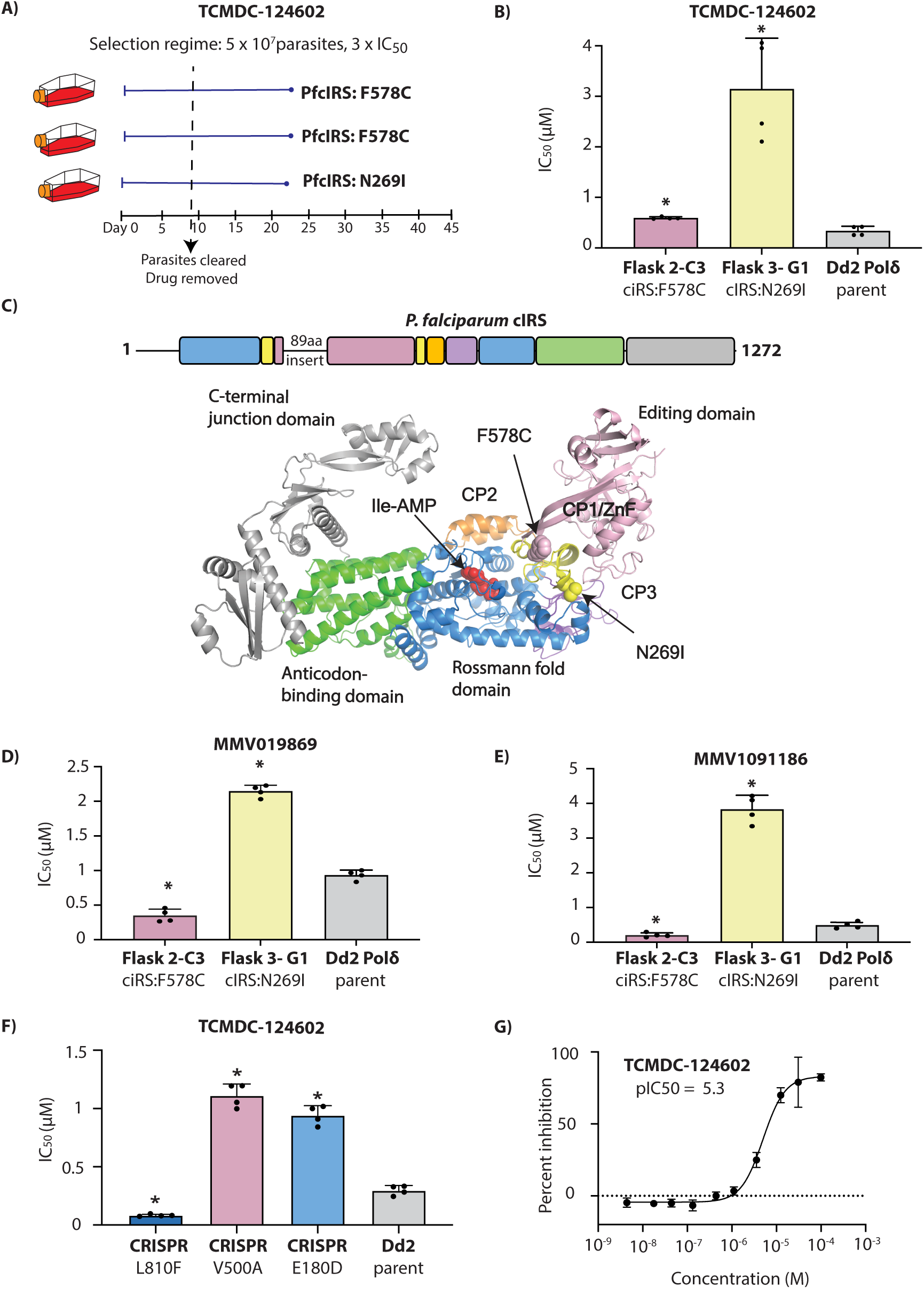
TCMDC-124602 targets *P. falciparum* cytoplasmic isoleucyl-tRNA synthetase (*Pf*cIRS). **(A)** Single-step resistance selection with TCMDC-124602 was performed in triplicate using 5 × 10⁷ Dd2-Polδ mutator parasites. Mutations in *Pf*cIRS were identified in all three selections. **(B)** Clonal lines from two independent flasks (Flask 2-C3 and Flask 3-G1) were tested for resistance to TCMDC-124602. The parental Dd2-Polδ line was included as a control. **(C)** Schematic representation of *Pf*cIRS architecture, coloured by domain, indicating the locations of resistance-associated mutations identified during compound selection. **(D–E)** Clones (Flask2-C3 and Flask3-G1) were further evaluated for cross-resistance to structurally related *Pf*cIRS inhibitors MMV019869 and MMV1091186. **(F)** CRISPR-edited parasite lines bearing the V500A and E180D mutations in *Pf*cIRS exhibited reduced sensitivity to TCMDC-124602, whereas the L810F mutant showed hypersensitivity. Dd2 parental line served as a control. **(G)** *In vitro* enzymatic assays confirmed inhibition of recombinant *Pf*cIRS by TCMDC-124602. In **(B,D-G)**, each dot represents a biological replicate (n = 4); bars indicate mean ± SD and statistical significance determined by Mann-Whitney *U* tests (*p<0.05).

In relation to cIRS domain structure, N269I is part of the CP1/ZnF domain. F578C is the last residue of the editing domain right before the second part of the ZnF domain (**Figure 3C**). This is similar to the previously obtained E180D and N269K mutations that resulted from selection with MMV1091186. In contrast, selections with MMV019869 yielded mutations within the CP1 editing domain (S288I, V500A, C502Y), and selections with thiaisoleucine resulted in an L810F mutant near the isoleucine binding pocket [18, 19].

When exposed to previously studied compounds, MMV019869 (**Figure 3D**) and MMV1091186 (**Figure 3E**), the N269I and F578C mutant clones showed differential responses. The clone carrying the *Pf*cIRS-N269I mutation (Flask 3) was found to be highly resistant to both compounds, whereas the clone harbouring the F578C mutation (Flask 2) showed modest but significant hypersensitivity to both compounds with IC_50_ shift ∼0.4-fold relative to the parental line.

Leveraging our previously generated CRISPR-edited lines with mutations in the different regions of *Pf*cIRS (E180D, V500A, and L810F; [18, 19]), we examined these for their response to TCMDC-124602. We observed that the ZnF/hinge region mutant E180D, as well as the CP1 editing domain mutant V500A, showed cross-resistance to TCMDC-124602 (**Figure 3F**). In contrast, the L810F mutant line was hypersensitive to TCMDC-124602, with an IC_50_ 0.3-fold lower compared to the parental line **(Figure 3G)**.

Finally, we assessed the inhibitory activity of TCMDC-124602 against recombinant *Pf*cIRS using an *in vitro* enzymatic assay based on AMP-Glo, which quantifies AMP generated by the enzyme. TCMDC-124602 effectively inhibited recombinant *Pf*cIRS, yielding an IC₅₀ of 6.59 µM, thereby confirming that *Pf*cIRS is the molecular target of TCMDC-124602 in *Plasmodium* (**Figure 3G, Supplementary Table 7**).

### Mutations in *P. falciparum* cGMP-dependent protein kinase (*Pf*PKG) confer resistance to compounds TCMDC-141334 and TCMDC-140674

Compounds TCMDC-141334 and TCMDC-140674 belong to a class of trisubstituted imidazole-based compounds (**Figure 1A**). Resistance evolution experiments yielded recrudescent parasites in 3 out of 3 cultures for TCMDC-141334 (**Figure 4A**), and 2 out of 3 cultures for TCMDC-140674 (**Figure 4C**). Clones isolated from these cultures were tested for resistance, with all showing resistance to their respective selection compounds (**Figure 4B, D**). TCMDC-141334-resistant clones exhibited a 2.2- to 4.1-fold increase in IC_50_ compared to the parental Dd2-Polδ line, while TCMDC-140674-resistant clones showed a higher fold change, ranging from 6.6 to 7.3 (**Figure 4B, D**).

**Figure 4.**
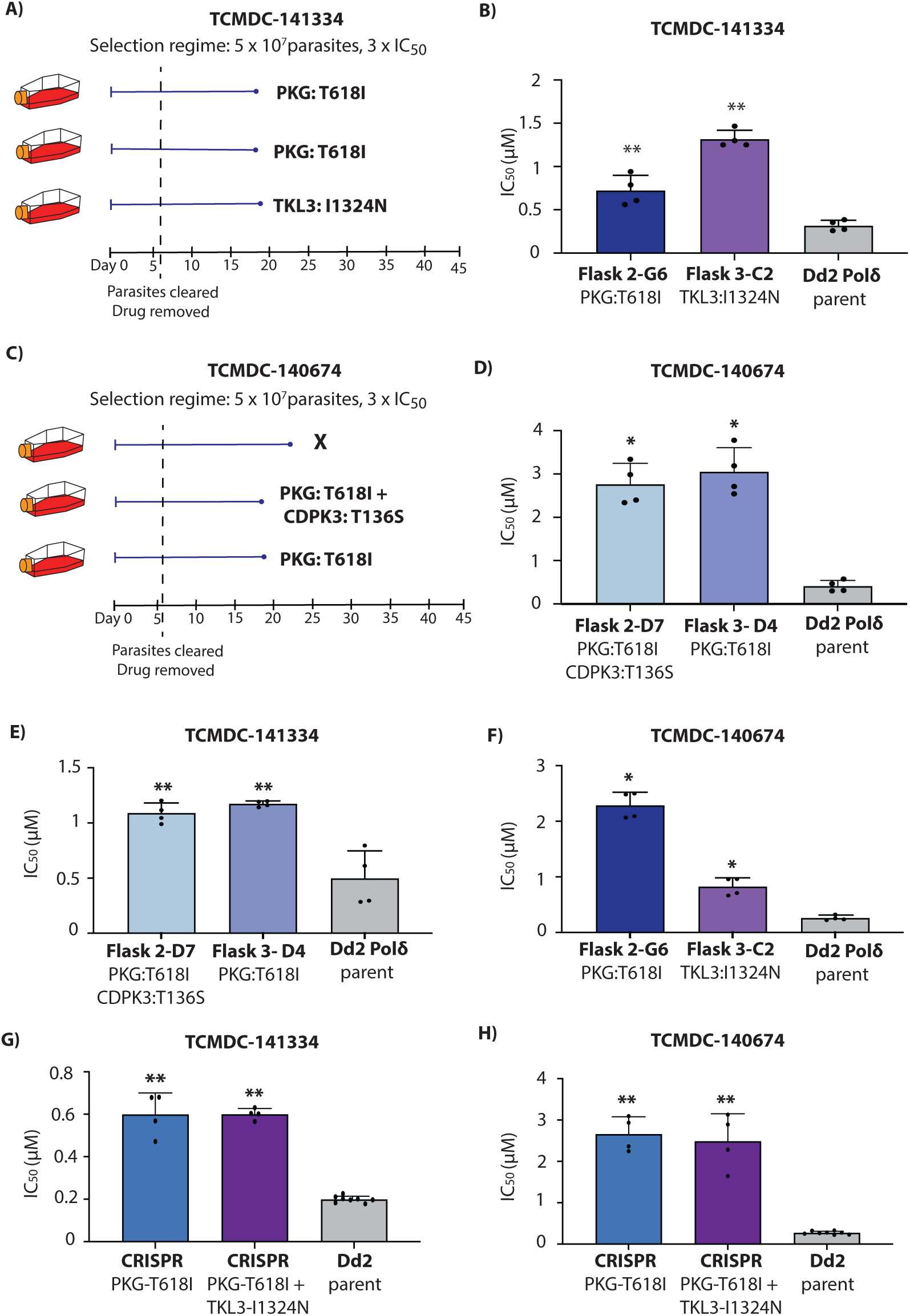
TCMDC-141334 and TCMDC-140674 target *P. falciparum* cGMP-dependent protein kinase (*Pf*PKG). **(A)** Single-step resistance selection with TCMDC-141334 was conducted in triplicate using 5 × 10⁷ Dd2-Polδ mutator parasites. Mutations in *Pf*PKG were identified in clones from two flasks, while a mutation in *Pf*TKL3 was observed in one flask. **(B)** Clonal lines from two independent flasks (Flask 2-G6 and Flask 3-C2) were evaluated for resistance to TCMDC-141334. The Dd2-Polδ parental line served as a control. **(C)** Resistance selection with TCMDC-140674, also performed in triplicate using 5 × 10⁷ Dd2-Polδ parasites, yielded *Pf*PKG mutations in clones from two flasks. One clone from Flask 2 additionally carried a mutation in *Pf*CDPK3. **(D)** Clones from Flask 2-D7 and Flask 3-D4 were tested for resistance to TCMDC-140674, with Dd2-Polδ used as a control. **(E–F)** Cross-resistance profiles of clones selected with TCMDC-140674 and TCMDC-141334 were tested against the reciprocal compounds. **(G–H)** CRISPR-edited parasite lines carrying the *Pf*PKG T618I mutation, as well as a double mutant line with *Pf*PKG T618I and *Pf*TKL3 I1342N, exhibited reduced sensitivity to both TCMDC-141334 and TCMDC-140674. The Dd2 parental line was included as a reference. In **(B,D-H)**, each dot represents a biological replicate (n = 4); bars indicate mean ± SD and statistical significance determined by Mann-Whitney *U* tests (*p<0.05, **p<0.01).

Whole genome sequencing of eight clones from the TCMDC-141334 selection and six clones from the TCMDC-140674 selection revealed mutations in a shared gene, cGMP-dependent protein kinase (*Pf*PKG, PF3D7_1436600) (**Figure 4A, C and Supplementary Table 8 and 9**). Furthermore, clones from both TCMDC-141334 and TCMDC-140674 selections resulted in the same mutation at the gatekeeper residue of *Pf*PKG (T618I) (**Supplementary Figure 1A**). Sequencing data also revealed MDR1 copy number amplification in the majority of clones analysed for both TCMDC-141334 and TCMDC-140674 (**Supplementary Table 4**).

In addition to mutations in *Pf*PKG, other mutations of potential interest were observed in a subset of sequenced clones. Resistance selections with TCMDC-141334 also identified three clones having no mutation in *Pf*PKG but a mutation (I1324N) in tyrosine kinase-like protein (*Pf*TKL3, PF3D7_1349300) (**Figure 4A, Supplementary Figure 1A and Supplementary Table 8**). The mutation identified in *Pf*TKL3 lies in the Sterile α-Motif (SAM) domain, considered essential for oligomerisation, protein-protein interactions, and is crucial for kinase activity [20] (**Supplementary Figure 1A**).

For resistance selections with TCMDC-140674, two clones shared a T136S mutation in calcium-dependent protein kinase 3 (*Pf*CDPK3, PF3D7_0310100) (**Supplementary Table 9**). This mutation was located in the kinase domain of *Pf*CDPK3. *Pf*CDPK3 is a prominent marker gene of mature *P. falciparum* ookinetes [21]. Its *P. berghei* ortholog is important for gliding motility in ookinetes [22] in response to a rise in cytosolic calcium, which in turn is controlled by PKG [23]. However, the expression of *Pf*CDPK3 is minimal during the blood stage and believed to play no important role during this stage of the parasite life cycle. While this mutation was therefore not considered to impact direct targeting of PfCDPK3 by TCMDC-140674, we note that interactions between mutations in PKG and a different member of the CDPK family affect asexual blood stage growth in *P. berghei in vivo* [24], raising the possibility of a compensatory mechanism for the observed PfCDPK3 mutation.

Parasites resistant to TCMDC-141334 exhibited cross-resistance to TCMDC-140674 (**Figure 4E and 4F**), and vice versa, consistent with them having similar targets and modes of action. To validate the role of drug-selected mutations in *Pf*PKG and *Pf*TKL3, we generated a CRISPR-edited mutant line introducing the T618I mutation in *Pf*PKG. The *Pf*PKG-T618I CRISPR-edited line showed a significant increase in IC_50_ for both TCMDC-141334 and TCMDC-140674, consistent with the resistance phenotype observed in the mutant parasites (**Figure 4G and Figure 4H**). To further explore the PfTKL3 SNV, we introduced the I1324N mutation into the *Pf*PKG-T618I edited parasites, generating double mutant parasites (*Pf*PKG-T618I and *Pf*TKL3-I1324N). The double mutant parasites exhibited no further significant shift in IC_50_ for either TCMDC-141334 or TCMDC-140674 relative to the single *Pf*PKG-T618I mutant (**Figure 4G and Figure 4H**). However, CRISPR introduction of the single PfTKL3-I1324N mutant in isolation resulted in a modest shift in IC_50_, confirming its relevance to the mode of action (**Supplementary Figure 1B**).

Molecular docking revealed different interactions of TCMDC-141334 and TCMDC-140674 with *Pf*PKG wildtype and *Pf*PKG-T618I (**Supplementary Figure 1C**). In *Pf*PKG wildtype, TCMDC-141334 is predicted to form 2 intermolecular hydrogen bonds, one with the N-H main chain between L620 and V621, and the other with C=O of the main chain between I681 and D682, whereas TCMDC-140674 also forms similar hydrogen bonds but one is with the side chain of T618 and the other with E625. When T618 was mutated to isoleucine, TCMDC-141334 was still able to form 2 intermolecular hydrogen bonds with the enzyme (one with N-H main chain between L640 and V621, and the other with the side chain of E625) but no intermolecular hydrogen bonding was detected between TCMDC-140674 and the mutant enzyme. This correlates with the higher IC_50_ shift in the *Pf*PKG-T618I parasite lines towards TCMDC-140674.

### Cross-resistance profiling of compounds that failed to generate resistance

*In vitro* evolution of resistance did not yield resistant parasites for the remaining 5 compounds (**Supplementary Table 1**). To further profile these compounds, we evaluated them using the AReBar (Antimalarial Barcode Sequencing) assay that comprised a pool of 52 barcoded parasites in the Dd2 or 3D7 strain background, that have mutations in genes recognized as antimalarial targets or resistance genes [25, 26] (**Supplementary Table 10**). As these mutant parasites are barcoded, their relative growth in the pool can be tracked by amplicon sequencing of each barcode, with outgrowth of specific lines offering insight into the mode of action of the query compound.

All five compounds (TCMDC-132158, TCMDC-142214, TCMDC-134122, TCMDC-137144, and TCMDC-133434) along with a control drug (DSM265) were tested at a concentration corresponding to 3 × IC_50_ of the Dd2 background (see **Figure 1B**), with assays conducted over a 14-day period. We found no growth of the pool for four of the compounds, while some growth was observed in the parasites exposed to TCMDC-134122 (**Figure 5A**). Nanopore sequencing of amplified barcodes was performed on the day 14 pool post-exposure for all five compounds. As anticipated, no parasite line was found to be enriched for the four no-growth compounds (**Figure 5B**). However, in the pool exposed to compound TCMDC-134122 we observed a modest enrichment above the 2.5 log2 fold change cutoff for three lines with the 3D7 background: 3D7-wild type, 3D7-MDR2-K840N, and 3D7-ACS10-M300I parasites (**Figure 5C, Supplementary Table 10**). To validate this observation, we performed dose-response assays with both the MDR2 and ACS10 mutant parasites; however, these showed no shift in IC_50_ relative to wild-type 3D7, indicating that they were false positives (**Figure 5D**). The overrepresentation of these parasites in the pool can be attributed to the fact that the IC_50_ for compound TCMDC-134122 is higher for 3D7 than for Dd2 (∼200 nM for Dd2 compared with ∼570 nM for 3D7; **Figure 1A**). Since the parasites were exposed to 3×IC_50_ of Dd2 (i.e. 648 nM) during the assay, this overall enrichment of 3D7 background lines (see triangle symbols in **Figure 5C**) may simply be due to the lower drug concentration used. Overall, these experiments suggest that the targets of these compounds are novel and not represented in the >30 modes of action captured in the AReBar pool.

**Figure 5.**
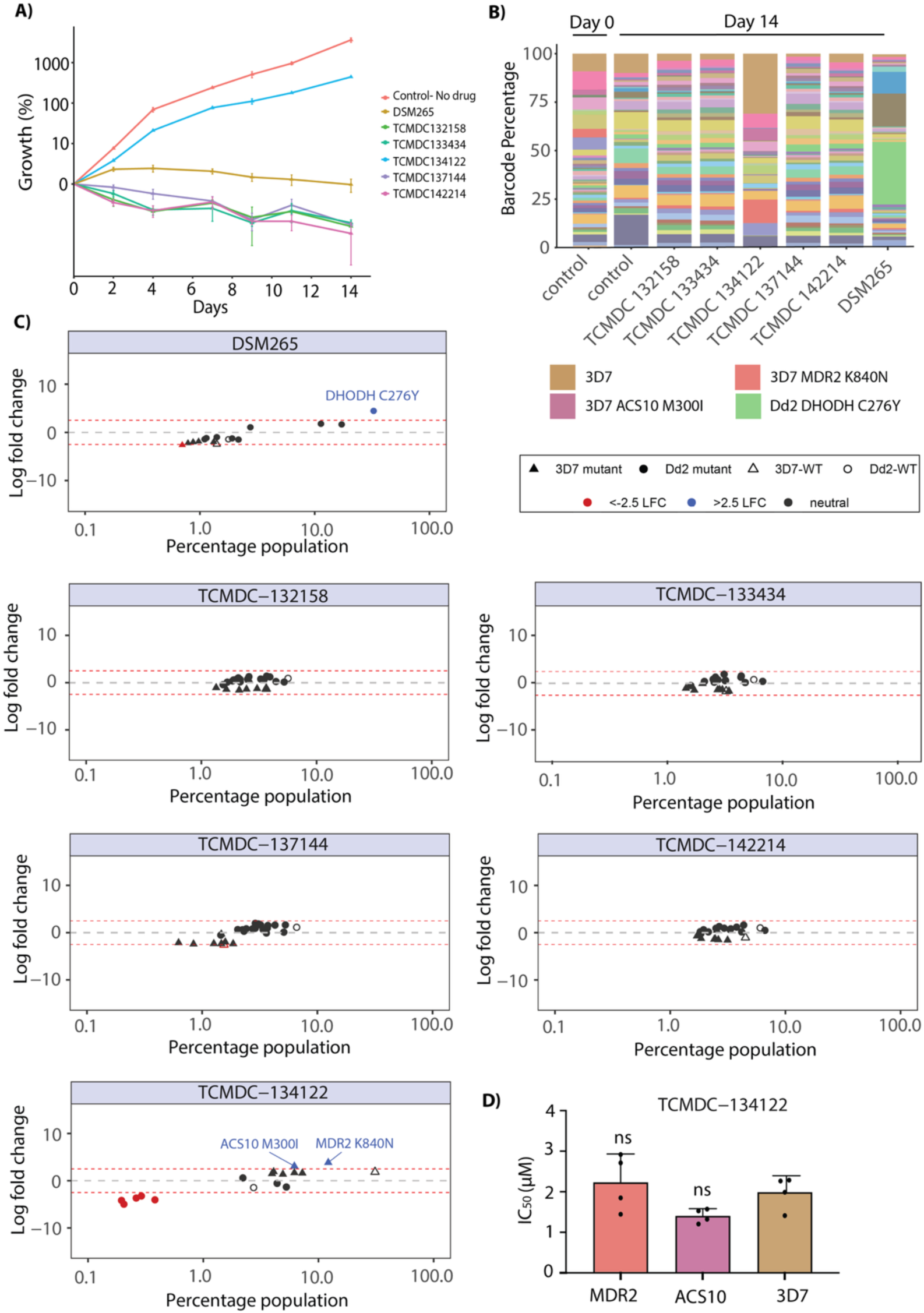
Cross-resistance profiling of non-resistant compounds using the AReBar (Antimalarial Barcode Sequencing) assay. **(A)** Growth curves for five compounds that previously failed to generate resistant parasites, alongside DSM265 (positive control with a known target) and a no-drug control. **(B)** Stacked bar plots showing the relative abundance of different barcoded parasite lines before and after 14 days of AReBar assay treatment with each compound. **(C)** Log₂ fold change (LFC) values for each barcode population following treatment with the five test compounds and DSM265, indicating potential compound-specific selection. Plots show LFC (y-axis) for individual lines in the pool, indicated by circles (Dd2 background) or triangles (3D7 background); dotted lines indicate 2.5 LFC. **(D)** Bar graph showing the drug response of MDR2 and ACS10 mutant cell lines (two candidate targets identified from AReBar analysis) following treatment with TCMDC-134122.

## Discussion

Drug resistance remains a significant obstacle to malaria eradication, highlighting the urgent need for novel antimalarials with distinct modes of action. Understanding the molecular targets of candidate compounds not only facilitates rational drug design and chemical optimization to enhance potency and binding affinity but also enables selectivity profiling to minimize off-target effects on human homologs. In this study, we investigated the mode of action of a panel of compounds previously reported to exhibit dual-stage activity against both blood- and liver-stage *Plasmodium* parasites. Using a mutator *P. falciparum* line, we performed *in vitro* resistance selections followed by whole-genome sequencing to identify potential targets. Resistance was successfully generated for four compounds—TCMDC-125075, TCMDC-124602, TCMDC-141334, and TCMDC-140674. Genomic analysis of resistant clones revealed mutations in *Pf*AcAS, *Pf*cIRS, and *Pf*PKG, implicating these proteins as putative drug targets. Despite multiple attempts, resistant parasites could not be selected for the remaining compounds, suggesting a lower propensity for resistance development.

*Pf*AcAS plays a vital role in generating acetyl-CoA within the nucleo-cytoplasmic compartment [17, 27]. In the nucleus, acetyl-CoA serves as a key substrate for the acetylation of histone lysine residues, an essential epigenetic modification that relaxes chromatin structure by weakening the interaction between DNA and nucleosomes. These acetylation marks are recognized by bromodomain-containing proteins, which facilitate the recruitment of transcriptional machinery, ultimately promoting gene expression [17, 27]. Inhibition of *Pf*AcAS leads to decreased histone acetylation levels, thereby reducing transcriptional activation of genes critical for parasite viability [17]. *In vitro* resistance selection using TCMDC-125075 resulted in the emergence of two mutations in *Pf*AcAS: G594V and T648M. The T648M mutation has also been previously reported following resistance selection with the structurally related compound MMV019721 [17]. CRISPR-engineered parasites carrying the *Pf*AcAS-T648M and *Pf*AcAS-A597V mutations confirmed *Pf*AcAS mutations as causal for resistance to TCMDC-125075. Target validation was confirmed by direct inhibition of the recombinant *Pf*AcAS enzyme.

The second target identified belongs to the aminoacyl-tRNA synthetase (aaRS) family, which plays a fundamental role in protein synthesis by catalysing the activation and attachment of specific amino acids to their cognate tRNAs. *Plasmodium* species encode approximately 36 distinct aaRS enzymes, making this protein family an attractive class of targets for antimalarial drug development [28]. Recently, Istvan *et al.* identified cytosolic isoleucyl-tRNA synthetase (cIRS) as a multistage antimalarial target [18, 19]. Their study demonstrated that selection with thienopyrimidine compounds led to resistance-associated mutations in *Pf*cIRS. Here, we observed two additional mutations in cIRS, with N269I related to the previously obtained N269K mutation, located in the CP1/ZnF domain. The novel F578C mutation, at the edge of the editing domain, provided only modest (∼1.7-fold) resistance to TCMDC-124602. In contrast, against the related thienopyrimidines MMV1091186 and MMV019869, these *Pf*cIRS-F578C parasites were hypersensitive, suggesting that protective effect of this mutation is highly context-specific. Direct inhibition of *Pf*cIRS activity by TCMDC-124602 was confirmed using the AMP-Glo assay with recombinant protein.

The final target we identified was cGMP-dependent protein kinase (PKG), mutations in which were identified from resistance selections with compounds TCMDC-141334 and TCMDC-140674. In *Plasmodium* parasites, PKG is a cGMP-binding protein that likely serves as the key mediator of cGMP signalling [29–32]. *Pf*PKG plays a crucial role in the tightly coordinated processes of RBC egress and invasion, regulating the release of the protease SUB1 from exonemes into the parasitophorous vacuole during egress, and discharge of AMA1 from micronemes during subsequent invasion [29–32]. Additionally, *Pf*PKG is required for gametocyte activation, ookinete motility, and regulates both invasion and exit from hepatocytes [29, 30]. Given its essential role at various stages of the parasite’s lifecycle, *Pf*PKG has long been considered a promising drug target [33–35]. Structural differences between *Pf*PKG and mammalian PKG, including at the gatekeeper residue of the ATP-binding pocket, allow for specific targeting of *Plasmodium* PKG. Resistance selections with TCMDC-141334 and TCMDC-140674 identified a mutation in the gatekeeper residue of *Pf*PKG from threonine to isoleucine that reduced the sensitivity of both compounds by 2-3 and 6-7-fold, respectively.

A previous study identified the trisubstituted imidazole compound MMV030084 as active against multiple stages of the *Plasmodium* life cycle, including asexual blood-stage parasites, by inhibiting merozoite egress, as well as male gametocyte exflagellation and sporozoite-mediated hepatocyte invasion [33]. This work established *P. falciparum Pf*PKG as the primary molecular target of MMV030084. However, resistance selections with MMV030084 did not yield mutations in *Pf*PKG but instead revealed that mutations in tyrosine kinase-like protein 3 (*Pf*TKL3) conferred low-level resistance. Although *Pf*TKL3 is a non-essential gene [33] and not the direct target of MMV030084, CRISPR-based introduction of the selected mutation confirmed its role in modulating sensitivity to the compound.

Baker *et al*. (2017) used Merck-derived imidazopyridine compound, ML1 (4-[7- [(dimethylamino)methyl]-2-(4-fluorophenyl)imidazo[1,2-a]pyridine-3-yl]pyrimidin-2-amine), as a chemical starting point to synthesize new analogs [34]. These were tested for inhibition of recombinant wild-type *Pf*PKG and a gatekeeper mutant (T618Q), to evaluate the role of the gatekeeper pocket in compound binding. Promising analogs were subsequently tested against *P. falciparum* asexual blood stages and further evaluated in the *Pf*PKG T618Q mutant parasite line to confirm on-target activity. Despite prolonged exposure of parasites to sub-lethal concentrations of these inhibitors, no resistant lines were recovered.

In the present study we report the emergence of a mutation in the *Pf*PKG gatekeeper residue through *in vitro* resistance selection using compounds targeting *Pf*PKG. This outcome was likely facilitated by the use of the Dd2-Polδ parasite line, which exhibits an elevated mutation rate, thereby enabling the evolution of resistance-conferring mutations in *Pf*PKG that were not observed in previous studies. Notably, for TCMDC-141334, we also observed a mutation in *Pf*TKL3, similarly to MMV030084. *Pf*TKL3 possess a kinase catalytic domain at the C-terminal region of the protein and contains a putative sterile alpha motif (SAM) domain which mediates oligomerisation and protein-protein interaction. Here we identified the I1324N mutation in *Pf*TKL3 in one flask of the TCMDC-141334 selection. This mutation is present in the SAM domain (amino acid positions 1306-1372), suggesting that it may hamper or modulate protein-protein interactions. Additionally, protein–protein interactions involving *Pf*TKL3 have been proposed to explain its role in modulating sensitivity to PKG inhibitors. One possibility is that *Pf*TKL3 mutations may alter phosphorylation patterns of one or more essential *Pf*PKG substrates, indirectly compensating for *Pf*PKG inhibition. However, the precise mechanism by which *Pf*TKL3 mutations confer resistance remains unclear and requires further investigation.

*In vitro* evolution of resistance has been a powerful approach for elucidating compound mode of action. Here we identified the targets for four of nine compounds, nonetheless drug resistance selections were unsuccessful for the remaining five compounds: TCMDC-132158, TCMDC-142214, TCMDC-134122, TCMDC-137144, and TCMDC-133434. To explore whether these compounds might be modulated by resistance mechanisms or have a characterized target, we adopted an alternative strategy using the AReBar assay, a pool of barcoded drug-resistant parasites with resistance mutations covering >30 modes of action. None of the compounds yielded validated leads through this assay, suggesting that they may act on previously unexplored drug targets, or potentially may hit multiple targets, making resistance generation challenging. Thus, they may warrant further investigation of their mode of action using orthogonal approaches such as affinity-based or thermal shift proteomics.

## MATERIALS AND METHODS

### Parasite cultivation and transfection

*P. falciparum* parasites were cultured with fresh human erythrocytes at 3% hematocrit in RPMI 1640 medium (Gibco), supplemented with 0.5% Albumax II (Gibco), 25 mM HEPES (Sigma), 1x GlutaMAX (Gibco), and 25 μg/mL gentamicin (Gibco). The following parasite strains were used: 3D7, Dd2 (MR4, MRA-156, T. Wellems), GB4 (MR4, MRA-925, T. Wellems), UG659 (P. Rosenthal [36]), Cam/PH0212C (R. Fairhurst [37]), and Cam3.II/ PH0306-C (R. Fairhurst [38]). The cultures were maintained under a controlled tri-gas atmosphere of 1% O_2_, 3% CO_2_, and 96% N_2_. Human erythrocytes were ethically sourced from healthy, anonymous donors through the Scottish National Blood Transfusion Service (SNBTS) or the National Health Services Blood and Transplant (NHSBT). The use of red blood cells was with approval from the University of Dundee Schools of Medicine and Life Sciences Research Ethics Committee, and the NHS Cambridgeshire Research Ethics Committee and Wellcome Sanger Institute Human Materials and Data Management Committee.

Ring-stage parasites were synchronized using 5% sorbitol, while late-stage parasites were synchronized with a 63% Percoll density gradient [39]. Transfections of ring-stage parasites were performed using the Gene Pulser Xcell electroporator (Bio-Rad). All gene sequences were retrieved from the PlasmoDB database (release 68) [40, 41].

To create the PKG and TKL3 mutants, the pDC2-Cas9-gRNA-donor plasmid was transfected into the *P. falciparum* Dd2 strain. For transfections, 50 µg of donor plasmid containing the sgRNA and a homology region of 810 base pairs for *pfpkg* and 578 base pairs for *pftkl3*, centred on the target mutation site was used. Guide RNAs for *pfpkg* (5’- CTAACAGAATTAGTAACAGG-3’ and 5’- CCTGTTACTAATTCTGTTAG-3’) and *pftkl3* (5’- GATATTTGCTAGAAGATTAAG-3’ and 5’- CTTAATCTTCTAGCAAATATC-3’) were designed using Benching (https://benchling.com/). Transfected parasites were selected using 5 nM WR99210 (Jacobus Pharmaceuticals) for 10 days, followed by a drug-free period until the parasites were detected. Clonal populations were then obtained by limiting dilution. Gene editing was validated using allele-specific PCR followed by Sanger sequencing with internal primers. For *pfpkg*, the primers used were: forward primer 5’- GAGCTTTATTATATGATGAACCAAGAACAGCTTCT-3’, reverse primer 5’-GGCCGGTTAATATATCACGGAAAATTTCT-3’, and the sequencing primer 5’- GTGTTAGTAAAAGAAGTATTATTAATTTAAATCAAC-3’. For *pftkl3*, the primers used were: forward primer 5’-AAGCCCCATGAATATTATTGTAACATC-3’, reverse primer 5’- ACTTGAGCATAGCTACCTCCTCC-3’, and the sequencing primer 5’- TGGAATAATACACGATTGATGGGTG-3’.

### *In vitro* drug resistance selections using Dd2-Polδ

The mutator parasite line Dd2-Polδ was used to generate resistance [16]. Parasites were exposed to 3×IC_50_ of the compound, and parasite viability was monitored daily using microscopy and Giemsa staining. After 5-10 days of treatment, no viable parasites were detected by blood smear. The compound was then removed, and the cultures were maintained by refreshing the media every other day, with fresh blood added weekly (by discarding half of the culture). Parasites re-emerged approximately three weeks after drug washout. For recrudescent cultures, compound susceptibility was assessed through dose-response assays. Clonal parasite populations were obtained by limiting dilution, and genomic DNA was extracted for whole genome sequencing.

### Compound dose-response assay

Compound dose-response assays were performed in flat-bottom 96-well plates. Tightly synchronized ring-stage parasites were adjusted to 1% parasitaemia at 1% haematocrit and exposed to a two-fold serial dilution of the compounds plated using Tecan D300e digital dispenser. As controls, the assay plate included untreated parasites and parasites treated with 700 nM DHA only. After 72 hours of incubation, parasite growth was measured by lysing the cells with a 2× lysis buffer containing 10 mM Tris-HCl, 5 mM EDTA, 0.1% w/v saponin, and 1% v/v Triton X-100, supplemented with 2× SYBR Green I (Molecular Probes). Fluorescence was recorded using a FluorStar Omega v5.11 plate reader. IC_50_ values were determined using GraphPad Prism v9, and statistical significance was assessed with a two-sided Mann-Whitney U test. Each assay was conducted in technical duplicate, with a minimum of three biological replicates, as described in the figure legends.

### AcAS Rapid-Fire QQQ Assays

Human Acetyl-CoA synthetase (HsAcAS) was assayed in 100 mM HEPES (pH7.4), 100 mM NaCl, 250 mM KCl, 50 mM MgCl2, 1 mM DTT, 0.005% NP40 substitute, 400 µM ATP, 70 µM acetate, 40 µM CoA and 2 nM recombinant HsAcAS. *P. falciparum* Acetyl-CoA synthetase (PfAcAS) was assayed in 100 mM HEPES (pH7.4), 100 mM NaCl, 250 mM KCl, 50 mM MgCl2, 1 mM DTT, 0.005% NP40 substitute, 150 µM ATP, 100 µM acetate, 20 µM CoA and 5 nM recombinant PfAcAS. Assay ready plates containing different concentrations of inhibitor were prepared using an Echo® 550 acoustic dispenser (Beckman) dispensing 150 nL of compound per well. Assays were conducted in a total volume of 15 µL at room temperature (30 minutes for HsAcAS, 50 minutes for PfAcAS) before being quenched with the addition of 85 µL 1% formic acid containing 0.5 µg/mL n-Propionyl coenzyme A (Sigma P5397) as an internal standard. Reaction products were detected using a RapidFire 365 system (Agilent, Santa Clara, CA) coupled with a triple quadrupole mass spectrometer 6740 (Agilent). The samples were loaded onto a D-hypercarb cartridge (Agilent) using deionized water at a flow rate of 1.5 mL/min and eluted to the mass spectrometer using 5 mM ammonium acetate in deionized water/acetonitrile/acetone (proportions of 2/1/1, v/v) at a flow rate of 1.25 mL/min. The sipper was washed to minimize carryover with 0.1% TFA in deionized water followed by 0.1 % TFA in acetonitrile/deionized water (95%/5%, v/v). Aspiration time, load/wash time, elution time, and re-equilibration time were set to 600, 3000, 5000, and 500 ms, respectively, with a cycle time of approximately 10s. The triple-quadrupole mass spectrometer with electrospray ion source was operated in positive multiple reaction monitoring (MRM) mode. The detailed setting for the mass spectrometer parameters was as follows: capillary voltage, 3000 V; gas temperature, 350°C; gas flow, 7 L/min; nebulizer, 40 psi; sheath gas temperature, 300°C’ sheath gas flow, 11L/min; and nozzle voltage, 1500 V. The MRM transitions for acetyl-CoA as a reaction product were set as 810.1/303.1 & 810.1/428.1 (quantifier & qualifier with dwell time of 50 ms, 100V fragmentor, and collision energies of 30 & 28 eV respectively) and for n-propionyl-CoA as an internal standard as 824.2/317.2 & 824.2/428.0 (quantifier & qualifier with dwell time of 50 ms, 100V fragmentor, and collision energies of 32 & 26 eV respectively). The mass resolution window for both parental and daughter ions was set at as a unit (0.7 Da). Peak areas were integrated, and area ratios of acetyl-CoA to the internal standard (n-propionyl CoA) were used for quantitation. Data were converted to % inhibition relative to the uninhibited enzyme control and fitted to a four-parameter logistic.

### Recombinant protein expression and purification

*Pf*AcAS, *Pf*AcAS-T648M, *Pf*AcAS-A597V, and *Hs*AcAS were expressed and purified as described previously [17]. The gene encoding *Pf*cIRS was codon optimised for expression in baculovirus and synthesised and subcloned into with a pTwist Kan vector (Twist Biosciences). The gene was subsequently cloned into a pFB-CT10HF-LIC by ligation independent cloning to append the full length expressed *Pf*cIRS (1-1273) with a C-terminal, TEV-cleavable His_10_-FLAG tag. The resulting clone was sequenced by DNA Sequencing and Services (MRC PPU, Dundee) to confirm identification of the clone. pFB-CT10HF-LIC was a gift from Nicola Burgess-Brown (Addgene plasmid # 39191; http://n2t.net/addgene:39191 ; RRID:Addgene_39191) The resulting plasmid was transformed into MAX Efficiency™ DH10Bac Competent Cells (Gibco) and screened by blue/white screening to confirm insertion. White colonies were grown in insect cell media prior to bacmid production. Bacmids were transfected into Sf9 cells conditioned in Sf900-iii SFM (Gibco) in 24-well plates seeding at 0.4×10^6^ cells using Insect GeneJuice Transfection Reagent (Sigma). P0 viruses were removed after 7 days incubation at 26.5 °C. Subsequent virus amplification was carried out from P0 (1 mL) to P2 (200 mL) in Sf900-iii SFM media in shaker flasks at 26.5 °C, 125 rpm, monitoring the cell viability and size. For expression, Sf9 cells were spilt to 1.5×10(^6^) cells in 500 mL volumes in Sf900-iii media in 2.5 L flasks (Nunc) and grown at 26.5 °C 125 rpm for 48 hours post virus infection, prior to centrifugation at 1000 x g for 10 minutes before being frozen until required. Cells were resuspended in lysis buffer (50 mM HEPES, 500 mM NaCl, 0.5 mM TCEP, 5 mM MgCl_2_, pH 8, DNase I, protease inhibitor cocktail) and lysed with a continuous flow cell disruptor (Constant Systems) at 30 kPSI. Protein was purified by Ni-NTA affinity chromatography (linear gradient 10-500 mM imidazole), His_10_-FLAG tag removal by TEV cleavage and reverse IMAC, and size exclusion chromatography. Protein was concentrated to 1.5 mg/mL in 25 mM HEPES, 500 mM NaCl, 0.5 mM TCEP, 5 mM MgCl_2_, pH 8 and stored at -80 °C.

### PfcIRS AMP-Glo™ Assays

*P. falciparum* isoleucyl tRNA synthetase (PfcIRS) was assayed in 50 mM HEPES (pH7.4), 100 mM NaCl, 30 mM KCl, 10 mM MgCl2, 1 mM TCEP, 0.1 mg/mL bovine serum albumin, 0.015% NP40 substitute, 5 µM ATP, 50 µM isoleucine, 0.025 mg/mL *in vitro* transcribed tRNAile and 5 nM recombinant PfIleRS. Assay ready plates containing different concentrations of inhibitor were prepared using an Echo® 550 acoustic dispenser (Beckman) dispensing 40 nL of compound per well. Assays were conducted in a total volume of 4 µL at room temperature for 180 minutes before being stopped with the addition of AMP-Glo™ reagents (Promega). Luminescence was quantitated using a Pherastar FX (BMG Labtech) and data converted to % inhibition relative to the uninhibited enzyme control before fitting to a four-parameter logistic.

### Modelling and computational docking

A homology model of PfcIRS was generated using AlphaFold [42, 43] while the structure of PfPKG was obtained from PDB (8EM8). The molecular structures of two compounds (TCMDC-141334 and TCMDC-140674) were prepared using Avogadro. The mutated residue in each enzyme was generated using PyMol version 2.5.2. DiffDock (https://build.nvidia.com/mit/diffdock), a diffusion generative model for molecular docking, was used to dock the compounds into both wildtype and mutant enzymes [44, 45]. DiffDock provides the affinity score of docked ligand poses, in which the first rank ligand was used for structural analysis. All structures were elucidated using PyMol version 2.5.2 [46].

### Whole genome sequencing analysis

Genomic DNA was extracted from resistant clones and Dd2-Polδ parent lines using DNeasy Blood & Tissue Kit (Qiagen). DNA libraries were generated using Nextera XT DNA Library Preparation Kit (Illumina), pooled, and sequenced on either Illumina NovaSeq 6000 (TCMDC-125075, TCMDC-141334, TCMDC-140674) or NovaSeq X Plus (TCMDC-124602) in the 100 bp paired end read (PE100) configuration. Raw sequencing reads were aligned to Plasmodium falciparum 3D7 reference genome (PlasmoDB v13.0) using BWA-MEM [47] and pre-processed following standard GATK protocols [48]. GATK HaplotypeCaller version 4.2.2.0 was used to call single nucleotide variants (SNVs) and indels, which were hard filtered based on the following exclusion criteria: ReadPosRankSum > 10 or < -10, Quality by Depth (QD) < 1.5, or Mapping Quality Rank Sum (MQRankSum) < -14. Variants were annotated using SnpEff [49] with a custom database built from the 3D7 reference GFF from PlasmoDB v13.0. To assess copy number variants (CNVs), genic coverage was standardized and denoised against a Dd2 panel of normals (as described in Luth et al. 2024, [50]) using GATK DenoiseReadCounts, yielding denoised log2 copy ratios per gene. Contiguous groups of genes with elevated log2 copy ratio were considered potential amplifications. Amplification CNVs were validated by the presence of discordant reads by manual inspection using IGV (Integrative Genomics Viewer) [51].

### Antimalarial resistome barcode sequencing (AReBar) cross-resistance assay

The AReBar assay was performed on a pool of 52 barcoded lines cover 33 modes of action, essentially as described in [25] (**Supplementary Table 10**). The lines possess resistance mutations in either targets or resistance genes, in both the Dd2 and 3D7 backgrounds (Supplementary Table 10), and were predominantly generated by the MalDA consortium [52, 53]. Each line was barcoded at the *pfpare* locus (PF3D7_0709700, [54]) with a distinct 11bp barcode sequence. Triplicate cultures of the pool (1ml volume) were challenged with compound, with query compounds tested at 3×IC_50_ over the 14 day assay period. Growth of the pool was monitored by flow cytometry (CytoFlex 5, Beckman) every 2-3 days, staining with 1×SYBR Green I, 0.2µM Mitotracker Deep Red, and parasitemia adjusted to a range between 0.3%-5% parasitemia. Samples were harvested at day 0, 4 and 14, and lysed with 0.05% saponin, washed twice with PBS, and resuspended in 30µL PBS prior to barcode amplification. Barcode amplicons were sequenced using a minION Mk1C (Oxford Nanopore), and counts analysed using DESeq2 [55] to quantify differentially represented barcodes relative to the no-drug control. Significance testing was performed by Wald test.

## Data availability

All associated sequence data are available at the European Nucleotide Archive under accession code PRJNA1289193

## Acknowledgements

We would like to thank Luma Godoy Magalhaes for help in rendering compound structures. This work was supported by funding to MCSL, OB and JCR from the Tres Cantos Open Lab Foundation (project TC266) and Wellcome (206194/Z/17/Z), and to MCSL from the Gates Foundation (INV-045096). MRL was supported in part by a Ruth L. Kirschstein Institutional National Research Award from the National Institute for General Medical Sciences (T32 GM008666). We also acknowledge support from the Structure-guided Drug Discovery Coalition funded by the Gates Foundation (INV-071352) and Wellcome funding for the Wellcome Centre for Anti-Infectives Research (203134/A/16/Z) to the University of Dundee. This publication includes data generated at UC San Diego IGM Genomics Center utilizing an Illumina NovaSeq 6000 that was purchased with funding from a National Institutes of Health SIG grant (#S10 OD026929).

## Author contributions

MR, NB and CS performed the *in vitro* drug selection experiments, drug sensitivity assays and parasite transfections, AP, JP, TE and DL performed the *in vitro* protein inhibition assays, JM and NB performed the structure analysis for cIRS and molecular docking modelling for PKG respectively. Whole-genome sequencing and data analysis were performed by MRL, DC and EAW. MR, NB, AP and MCSL planned the experiments and BB, EAW, JCR, OB, SMC, FJG and MCSL supervised the study. JCR, OB and MCSL contributed to funding acquisition. All authors contributed to writing the paper.

---

**Supplementary Figure 1.**
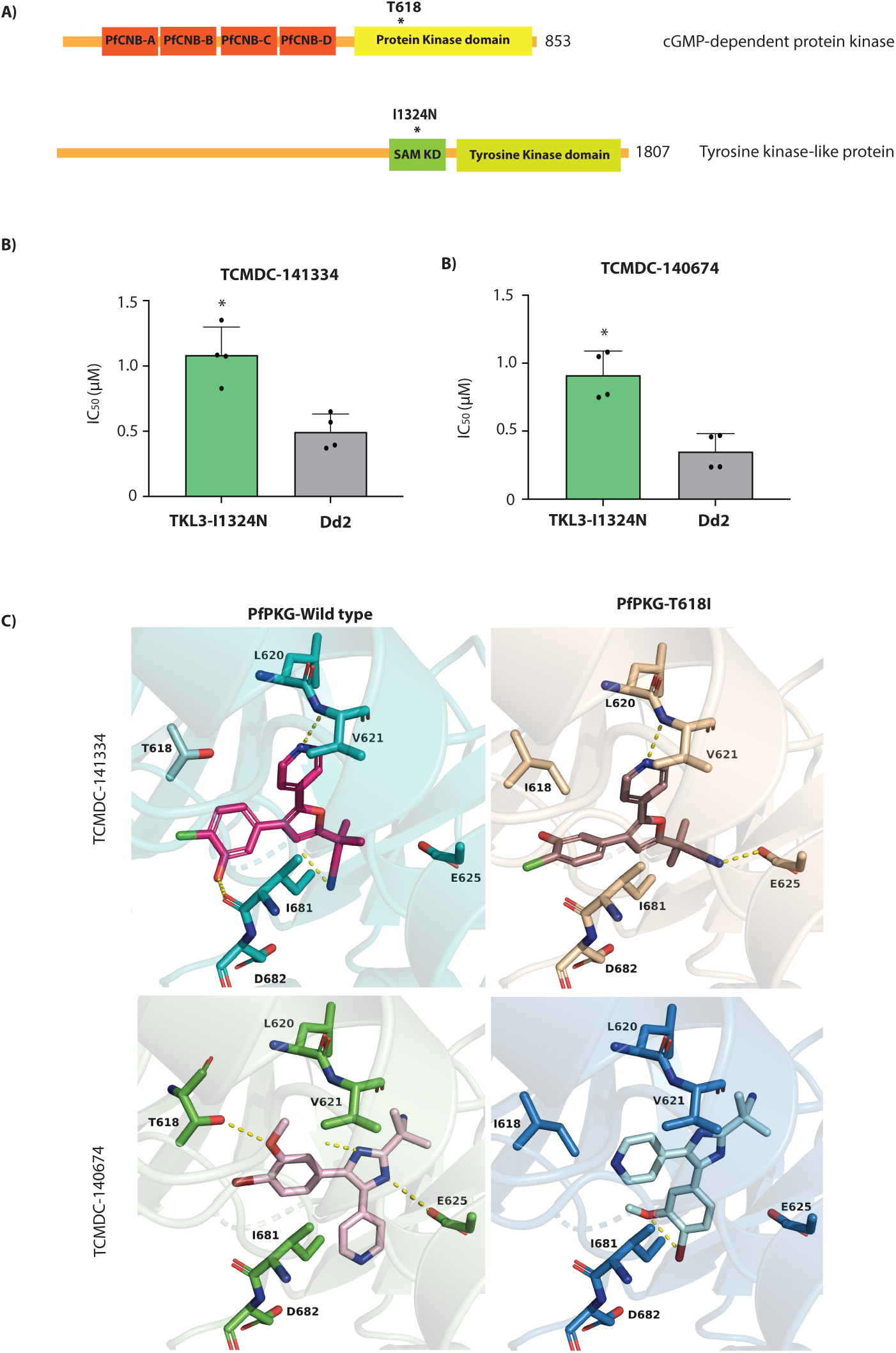
TCMDC-141334 and TCMDC-140674 target *P. falciparum* cGMP-dependent protein kinase (PfPKG). **(A)** Schematic representation of the domain architecture of PfPKG and PfTKL3. (**B)** CRISPR-edited parasite lines carrying the *Pf*TKL3 I1342N mutation exhibited reduced sensitivity to both TCMDC-141334 and TCMDC-140674. The Dd2 parental line was included as a reference. Each dot represents a biological replicate (n = 4); bars indicate mean ± SD and statistical significance determined by Mann-Whitney *U* tests (*p<0.05, **p<0.01). **(C)** Molecular docking models showing the binding of TCMDC-141334 and TCMDC-140674 to both wild-type PfPKG and the PfPKG-T618I mutant.

